# The transposable elements syndrome of wheat domestication

**DOI:** 10.64898/2026.01.28.702265

**Authors:** Guy Ben Zvi, Sariel Hübner

## Abstract

Domestication marks a turning point in human history, driving substantial genomic modifications in cultivated plants. Tetraploid wheat was one of the Neolithic founder crops and remains a major cultivated species today. We investigated how domestication altered the abundance and distribution of transposable elements (TEs) and structural variation (SV) in wheat, revealing genomic changes associated with key domestication traits. Our results demonstrate that extensive TE proliferation before and during the Pleistocene–Holocene transition expanded standing genetic variation and phenotypic diversity. This expansion has provided the raw material for selection by the early farmers including a Gypsy insertion in the *BTR1*-3B gene causing a loss of function and contributing to the establishment of the non-shattering phenotype. We further show that purifying selection was more efficient in purging TEs among wild populations, whereas domesticated wheat has maintained TE clusters around genes that are under selection. We propose a model in which climatic instability triggered genome-wide TE bursts, expanding genetic variation including at domestication traits. As climate stabilized, purifying selection gradually removed deleterious TEs insertions, while early farmers selectively preserved advantageous phenotypes, thus maintaining TE-rich regions around key domestication genes. This model provides an integrative framework linking climate driven genomic changes, selection, divergence and domestication.

## Introduction

Domestication marks a turning point in the history of humans and their surrounding ecosystem. The earliest evidence for domestication was excavated in the Fertile Crescent, an ecologically diverse region which lays between continents and a transition zone from desert to temperate climates (Zohary et al. 2012; Harlan and Zohary 1966; Bar-Yosef 1998; Frumkin et al. 2011). Owing to its geographic position, this region has been repeatedly shaped by climatic fluctuations throughout history, resulting in dramatic landscape transformations (Bar-Matthews et al. 1999). These processes continue today, with accelerating desertification driven by rising global temperatures (Lelieveld et al. 2012; IPCC 2023). Among the crops domesticated in the Fertile Crescent around 10,000 years ago, wheat has had the greatest global impact on human diets and is now cultivated across more than 200 million hectares worldwide (Shewry and Hey 2015; FAO 2024).

The domestication of the tetraploid wild emmer wheat (*Triticum turgidum* ssp. *dicoccoides*; WEW) has resulted with the emergence of the domesticated emmer wheat (*T. turgidum* ssp. *dicoccum*, DEW), and durum wheat (*T. turgidum* ssp. *durum*, DW). The hexaploid bread wheat (*T. aestivum*) arose later through hybridization between cultivated tetraploids and the diploid *T. tauschii* (DD genome) (Dvorak et al. 1998; Feldman et al. 2001; Luo et al. 2017). Despite the extensive research dedicated to elucidating the emergence of domesticated crops and specifically wheat, few controversial perspectives on the transition from a wild species into a cultivated crop remain (Abbo et al. 2014; 2021; Allaby et al. 2022; Purugganan 2019). The two leading models of plant domestication differ in their view on the tempo, scale, and intensity of selection. The protracted domestication model supports a slow evolutionary process over millennia across large, interconnected populations, driven by weak, unconscious selection (Allaby et al. 2008; Purugganan and Fuller 2009; Fuller et al. 2014). In contrast, the rapid domestication model posits that key domestication traits like the non-brittle rachis in cereals were fixed rapidly in localized founder events, while subsequent crop improvement traits evolved later (Harlan et al. 1973; Abbo et al. 2014). Empirical studies consistently showed that domestication has reduced genetic diversity, although the magnitude of this bottleneck remains debated and often reflects contrasting narratives of rapid versus protracted domestication (Haudry et al. 2007). Although domestication involved genetic bottlenecks that reduced overall diversity, many key domestication alleles likely arose from standing genetic variation predating cultivation (Studer et al. 2011). Most comparative analyses have focused on shifts in diversity and recombination between wild and domesticated taxa by looking at point mutations, particularly single-nucleotide polymorphisms (SNPs), which are now highly accessible for genomics studies. Recently, a comprehensive whole genome sequence data has become available for a tetraploid wheat collection allowing to locate the origin of domesticated wheat in Southeast Turkey and identify ancient contribution through introgressions from wild emmer populations in the southern Levant enriching the genetic makeup of domesticated wheat (Lev-Mirom et al. accepted). Nevertheless, domestication is not limited to shifts in point mutations frequencies and introgressions, and represents a broader genetic remodeling, producing a considerable genomic change and a suite of new traits referred as the domestication syndrome. These traits frequently arise from altered gene regulation or loss-of-function mutations in developmental regulators, often mediated by transposable element (TE) insertions (Doebley et al. 1995; Meyer and Purugganan 2013; Stitzer and Ross-Ibarra 2018). However, major genomic changes, namely structural variations (SVs) and large-scale rearrangements have been largely overlooked in domestication studies, despite their pervasive influence on genome evolution. This gap is especially pronounced in wheat, where population-scale analyses of TE dynamics remain scarce due to the technical challenges posed by its large, repetitive, and polyploid genome. Several cases illustrate the importance of regulatory and TE-mediated mutations in domestication. For example, the *Hopscotch* retrotransposon insertion upstream of *tb1* increasing apical dominance in maize (Doebley et al. 1995; Studer et al. 2011), the reduced *fw2*.*2* expression which enlarged tomato fruit size (Frary et al. 2000), and Mendel’s wrinkled pea phenotype caused by an Ac/Ds-like TE disrupting a starch-branching enzyme gene (Bhattacharyya et al. 1990).

Transposable elements (TEs) likely play a central role in domestication but remain difficult to explore due to challenges in detecting TE polymorphisms (Goubert 2023; Stitzer and Ross-Ibarra 2018). Among the different TE classes, long terminal repeat retrotransposons (LTR-RTs) and particularly the *Gypsy* and *Copia* superfamilies, dominate the wheat genome, thus these superfamilies are more frequent and therefore detectable even with short-reads sequence data. Recent advances in bioinformatic tools now facilitate the reliable detection of TE polymorphisms, enabling insights into their contribution to structural variation and crop evolution (Goubert et al. 2015; Zhou et al. 2020; Wicker et al. 2018).

To elucidate the role of TEs in wheat domestication, we investigate whole genome sequence data from 159 tetraploid wheat accessions representing WEW, DEW, and DW. Using complementary approaches, we characterized TE abundance and distribution along the genome of all samples, elucidating their contribution to population differentiation and signatures of selection. Our findings highlight key role of TEs in wheat domestication and provide a new molecular perspective on domestication and the emergence of crop species.

## Results

### Wheat domestication has shifted transposable elements abundance

To characterize the abundance of TE families and their distribution along the genome, we re-annotated the repetitive fraction of both wild emmer (“Zavitan”) and domesticated durum (“Svevo”) wheat reference genomes. The overall TE family composition was broadly conserved between genomes (Fig. S1). The most prevalent class was the long terminal repeat (LTR) retrotransposon *Gypsy*, accounting for approximately 40% of the genome, followed by *Copia* (14%), the terminal inverted repeat (TIR) DNA transposon family CACTA (8%), and all remaining TE families combined (5%). As expected, *Gypsy* elements were preferentially enriched in pericentromeric regions, whereas *Copia* and CACTA elements were primarily distributed along the chromosomal arms (Fig. 1a, Fig. S2).

**Figure 1.**
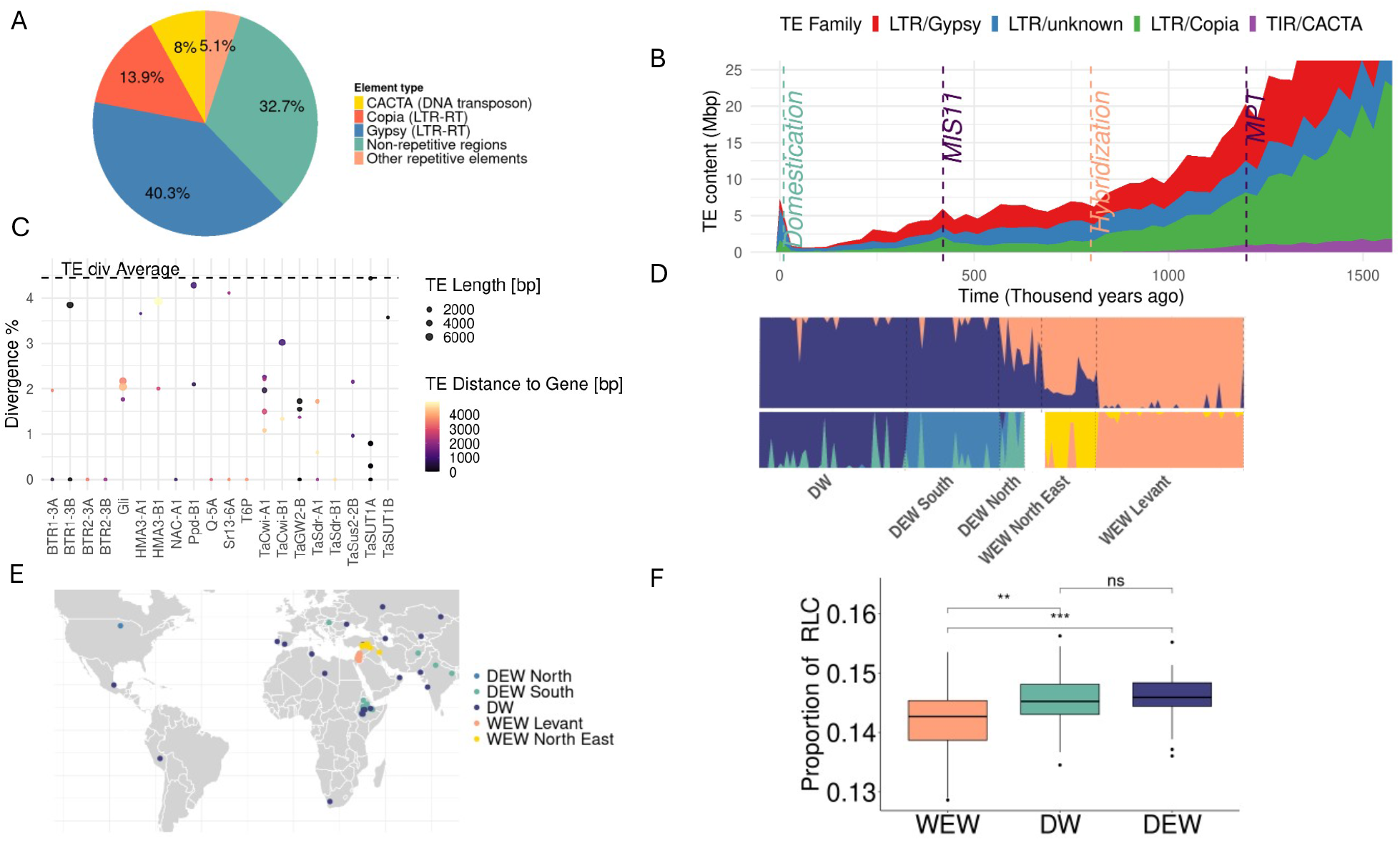
Exploring transposable elements (TE) abundance among wheat groups. **A)** Pie chart of TEs proportion in the genome. **B)** TE divergence accumulation along time in the *Svevo* genome. The x-axis represents the calculated time based on sequence divergence of proliferation events in genome, and the y-axis shows the intensity. Each color corresponds to a different TE family. Major climatic turnover events (MPT and MIS11), the tetraploid wheat ancestor’s hybridization and domestication are marked with vertical dashed lines. **C)** Divergence and characteristics of TEs associated with candidate genes. The plot displays the level of TE divergence (y-axes), distance for the gene (color gradient) and the size of TEs in the Svevo genome. The horizontal dashed line represents the average divergence of the TEs along the genome. **D)** Structure analysis of tetraploid wheat groups. The lower panel corresponds to the wild emmer (right) and domesticated subgroups (left), and the upper panel represents the assignment to specific wild or domesticated types among all accessions. **E)** A map showing the origin of each sample in the study colored by type. **F)** Comparison of *LTR/Copia* elements (RLC) proportion between the main groups where statistical significance for each comparison is indicated at the top.

Estimates of sequence divergence within TE families revealed a continuous accumulation of TEs over the evolutionary history of wheat, with notable bursts of TE proliferation. Scaling sequence divergence to estimated time frame was conducted using a fixed mutation rate of 1.67×10^-8^ and highlighted a notable TE proliferation during the transition from the Pliocene to the Pleistocene (Fig. S3). At the fine scale, small bursts of TEs were identified around the Mid-Pleistocene Transition (MPT) circa 1.2Mya, a blip burst right after the hybridization between the ancestor species or tetraploid wheat circa 0.8Mya, and a more pronounced burst 0.4Mya around the interglacial warm period of the Marine Isotope Stage 11 (MIS11). The signature of these TE bursts was detected in both wild and domesticated genomes confirming these events predated domestication. Notably, additional signal of TEs with very low divergence was also detected in both reference genomes which is likely attributed to the transition to the Holocene just before the domestication of wheat (Fig. 1b, Fig. S3). To further explore TEs in the Svevo genome that are characterized with low divergence, we calculated the sequence divergence of identified TEs relative to their closest copies in the genome. Overall, TE pair-wise divergence across the genome is characterized with a broad distribution (Fig. S4). However, several outliers with exceptionally low divergence were identified, indicating their recent spread along the genome. Among the recent TEs insertions we identified several TEs that were located closely or within key domestication genes, including BTR1-3B/3B, BTR2-3B/3B, Q-5A, NAC-A1, Ppd-B1 and other genes that are associated with the crop evolution (Fig. 1c).

To further quantify the differences in TE abundance among wild and domesticated wheats, we analyzed a diversity panel comprising 159 tetraploid wheat accessions (Table S1), including wild emmer wheat (*Triticum turgidum ssp. dicoccoides*; WEW), domesticated emmer wheat (*T. turgidum ssp. dicoccum*; DEW), and durum wheat (*T. turgidum ssp. durum*; DW). Genotype data was pruned for linkage disequilibrium (LD > 0.2), reducing the dataset to 2,656,229 SNPs for population structure analyses. Both fastSTRUCTURE and PCA supported the classification of accessions into three genetically distinct groups corresponding to WEW, DEW, and DW (Fig. 1c-d, Fig. S5). To identify potential misclassification or admixture events between wild and domesticated wheat, we tested the assignment of individuals to their expected group using K = 2. Nine accessions showed less than 50% assignment to their expected group and were removed from further analysis. Within the domesticated group, structure analysis revealed three subgroups corresponding to DW, southern DEW (primarily Ethiopian), and northern DEW (Eurasian). Among domesticated accessions, eight were misassigned to their expected group and were excluded. Consequently, only six accessions were confidently assigned to the northern DEW group. Due to genetic differences between the northern and southern DEW these two subgroups could not be integrated into one group. Because the northern DEW group was too small (n = 6) to stand robustly as a separate group it was excluded from further analyses. The final dataset comprised of 150 accessions of which 64 WEW, 28 southern DEW, and 48 DW (Fig. 1d-e, Table S1) with significant reduction in genetic diversity along domestication gradient (Fig. S6).

To quantify the proportion of TEs in each group, we used a reference-free pipeline which performs *de novo* assembly from a representative sample of sequences for each accession separately. To identify the number of reads that properly represent the proportion of TEs in the genome, we calibrated the pipeline by sampling an increasing number of reads from 0.5 to 20 million. The proportion of TEs identified has remained stable even at the sub-class level after 10M reads, thus indicating that sampling has been saturated (Fig. S7). Therefore, we fixed sampling at 10M reads for each accession across the entire collection. Expectedly, LTRs employed most of the TE fraction along the genome, of which the *Gypsy* family was the most abundant among accessions of all three groups (Fig. S8). The second most abundant LTR family correspond to *Copia* which was significantly more abundant (F = 9.068, p = 2.07 × 10^--−4^) among domesticated accessions than in the wild. Higher rate of *Copia* was observed in both DEW and DW compared to WEW, thus indicating that this trend is associated with the transition from a wild species into a domesticated crop (Fig. 1f).

Based on the profile of sequence divergence and comparison of TEs content and variation, we hypothesize that following the TE bursts, insertions were gradually purged by natural selection over time, removing deleterious elements. Domestication led to less efficient purging due to the combined effects of the bottleneck and genomic fixation, while TEs continued to be efficiently removed in wild populations. To test this, TE insertions were identified using the Zavitan as the reference. For comparison, the same analysis was conducted with Svevo as a reference, but the number of insertions was much lower (mean = 7,115) than identified with Zavitan (mean = 19,571). This is expected, as the domesticated Svevo contains more TEs that were purged in the wild which are not detectable when calling insertions against a reference. Analyzing the identified insertions, we generated a site frequency spectrum (SFS) for wild and domesticated groups. The wild SFS was enriched with rare variants and poor in high-frequency variants compared to the domesticated SFS, implying more efficient TE purging in the wild (Fig. S10). Since the profile of TE sequence divergence was similar between the wild and domesticated types prior to domestication, it is likely that domestication fixed and maintained a higher proportion of TEs that continued to be purged by natural selection in the wild.

### TEs are more abundant around domestication genomic hotspots

To investigate the impact of domestication on the genomic distribution of TEs, we compared TE occurrence between wild and domesticated wheat accessions. Sequencing reads from each accession were aligned to the Svevo reference genome, and percent coverage of each annotated TE was calculated. The Svevo reference was selected because many domestication-related TE insertions are absent in the wild reference, and the Svevo genome has a higher TE content, particularly of the *Copia* superfamily which was identified to be more abundant among domesticated accessions in the reference-free approach (Fig. 1f, Fig. 2a, Fig. S1).

**Figure 2.**
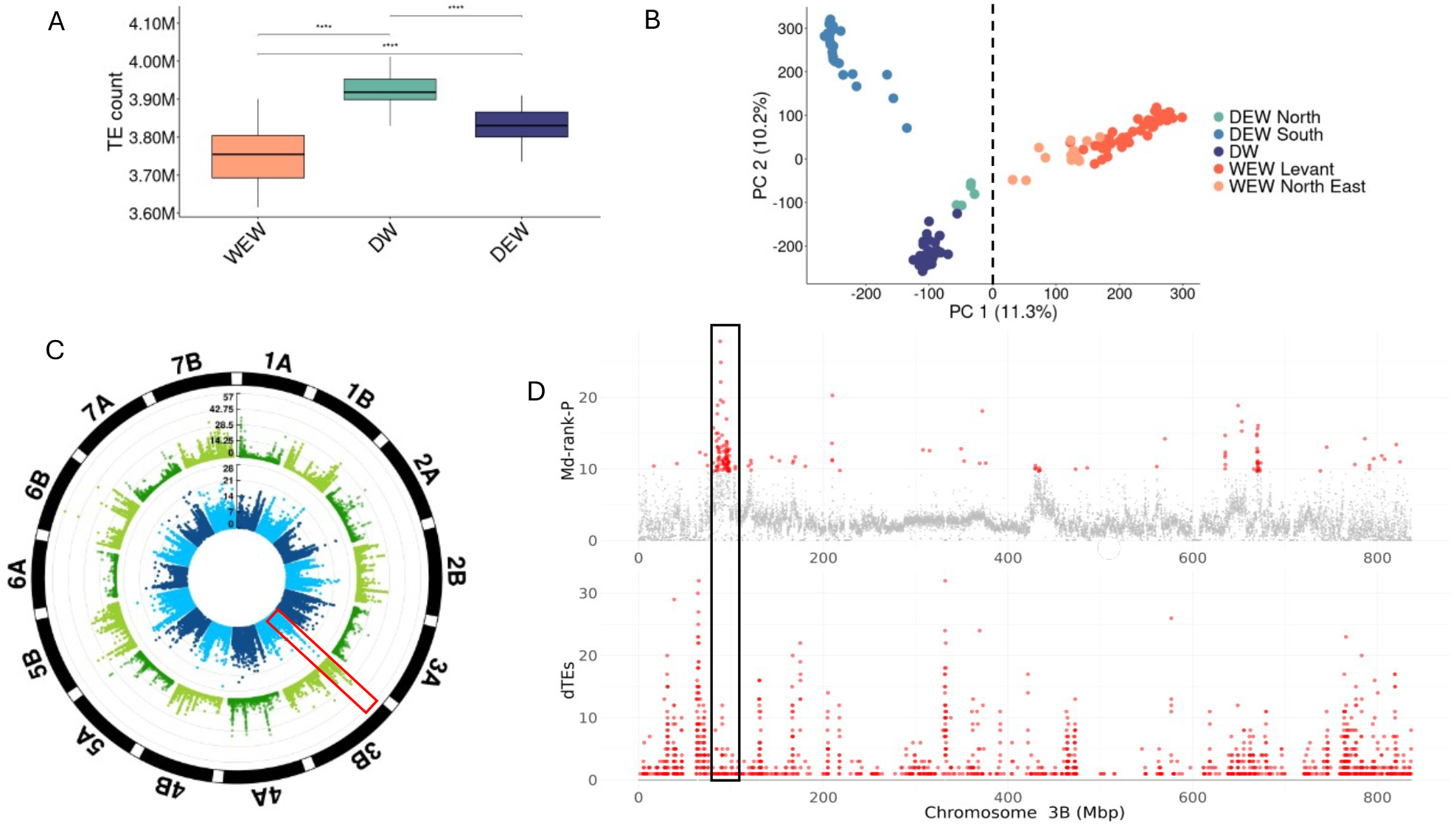
Identification of domestication related TE hotspots. **A**) Comparison of TE counts obtained from the occurrence of TEs is each accession. Statistical significance between groups is indicated at the top. **B**) Principal component analysis (PCA) based on the PAV matrix of TEs occurrence. Accessions are colored based on their assignment to a group and horizontal line splits between domesticated and wild TEs. **C**) Circular genome plot of genome scans. In the inner track depicts the selection test (Md-rank-P scores per window), and in the outer track corresponds to the dTEs counts in 50Kbp windows.A zoom in on the red frame indicated in (C) where the *btr1-3B* gene is marked with vertical dashed lines. The top panel corresponds to the selection analysis, and the bottom panel corresponds to top dTEs counts, both in 50Kbp windows.

To assess the robustness of this analysis and exclude potential reference biases, we replicated the analysis using the Zavitan reference genome (Fig. S11). Expectedly, less TEs were identified using the Zavitan as a reference, yet the overall trend was consistent in both reference genomes, thus indicating an increased TE content among domesticated accessions, supporting the reference-free analysis (Fig. 2a). Based on read coverage, TEs were classified as present (≥75% coverage), absent (0% coverage), or unknown (intermediate values). This conservative approach prioritizes lower false positives, which are more concerning for repetitive TE sequences, at the cost of potentially increased false negatives (Fig. S12). The TEs occurrences were compiled into a binary presence/absence variation (PAV) matrix for all accessions and used for downstream population-level comparisons. Overall, domesticated accessions exhibited a significantly higher number of TEs than wild accessions, confirming our previous results (Fig. 1f, Fig. 2a, Fig. S9, Fig. S11). To maintain higher confidence in identifying trends, further comparisons were made at the population level rather than for specific individuals.

To identify TEs that differentiate wild and domesticated wheat, a PCA was conducted using the PAV matrix of TE occurrences. The first principal component (PC1) explained 11.3% of the total variance and clearly separated wild from domesticated accessions (Fig. 2b). Interestingly, the differentiation based on TEs highlighted the north-eastern WEW population as the closest population to domesticated wheat, thus supporting the region of nowadays south-east Turkey and northern Syria as the origin of initial domesticated wheat form. The second component (PC2), accounting for 10.2% of the variance, further distinguished between domesticated emmer wheat (DEW) and durum wheat (DW) indicating that these two groups diverged also in their TEs landscape. Next, we extracted a list of TEs contributing to the wild-domesticated differentiation from PC1 loadings. TEs with strongly negative loading scores were classified as domesticated-associated TEs (dTEs), and TEs with strongly positive loadings were considered wild-associated TEs (wTEs). The top 1% of TEs from each tail of the loading distribution were selected for further analysis, yielding approximately 80,000 TEs (Fig. S13). To explore the genomic distribution of dTEs, we quantified dTE abundance within 50Kbp windows and targeted ‘dTE hotspots’. Overall, dTEs were clustered in 5% of the windows, compared to a random subset of TEs which were distributed along 20% of the genome. This indicates that dTEs tend to cluster within more specific genomic regions (Fig. S14).

To determine whether dTEs hotspots are located within genomic regions that were under selection during domestication, a genome scan analysis for footprints of selection was conducted in sliding windows. We calculated the differentiation index (F_ST_) between wild and all domesticated accessions, the strength of reduction in nucleotide diversity (π_D_/ π_W_), the extent of reduction in recombination rate (ρ_*D*_/ρ_W_), and the Tajima’s D score for the domesticated group (Fig. S15). All four statistics were then combined into a composite measure of selection by transforming the scores to a negative log rank-based p-values integrated with the Mahalanobis distance (Md-rank-P). We used this approach to avoid LD-based statistics which are biased by the integration of DEW and DW into one domesticated group and northern and southern WEW into one wild group. The top 5% of windows were considered as regions that were targeted by selection during domestication (Fig. S16). Notably, genomic regions that were targeted by selection were also consistently enriched with dTEs (t_Welch_ = 2.08, *p* = 0.03) and depleted with wTEs (t_Welch_ = -20.16, *p* < 2.2×10^-16^) compared to neutral genomic regions (Fig. S18). Overall, 9965 windows were identified as directly affected by domestication, of which 923 windows were also identified as dTEs hotspots (Fig. 2c, Table S2, Fig. S17). Within these regions, several key domestication related genes were identified including the major seed shattering genes *BTR1-3A, BTR2-3A, BTR1-3B, BTR2-3B*, the key domestication *Q* gene, and the vernalization gene *VRN-B3* (Table S3).

Interestingly, one of the top signals of selection and accumulation of dTEs was observed around the *BTR1-3B*, which is a key gene in the suppression of seed shattering in domesticated wheat. We found *Gypsy* dTEs identified exclusively among domesticated accessions and absent in the WEW. These dTEs are located within the 4Kbp insertion in the Svevo reference genome which was previously reported as causative for the gene loss of function and the non-brittle spike phenotype. Exploring this region in the recently assembled Langdon reference genome (Chen et al. 2025) supported the Gypsy insertion, yet this insertion spans over 20Kbp rather than 4Kbp as indicated in the Svevo genome (Fig. S20). This example underscores the potential role of TE insertions in the evolution and fixation of domestication-related alleles in wheat.

### Domestication TEs are associated with structural variation

To further elaborate the genomic divergence between wild and domesticated wheat, we compared the Svevo and Zavitan reference genomes using SyRI to identify structural variation (SVs) including duplications, inversions, and translocations. These SVs were filtered to remove instances where the SV is a TE, thus a total of 43,801 SVs were further analyzed. The largest SV variant detected was an inversion of 64,124,654 bp on chromosome 2A. Along the entire genome, SVs covered circa 20% of the Svevo and Zavitan genomes, with even distribution across chromosomes (Fig. 3a). Among SVs, we detected 608 large inversions, 5,738 translocation, and 18,159 duplications (15,807 in Svevo and 2352 in Zavitan).

**Figure 3.**
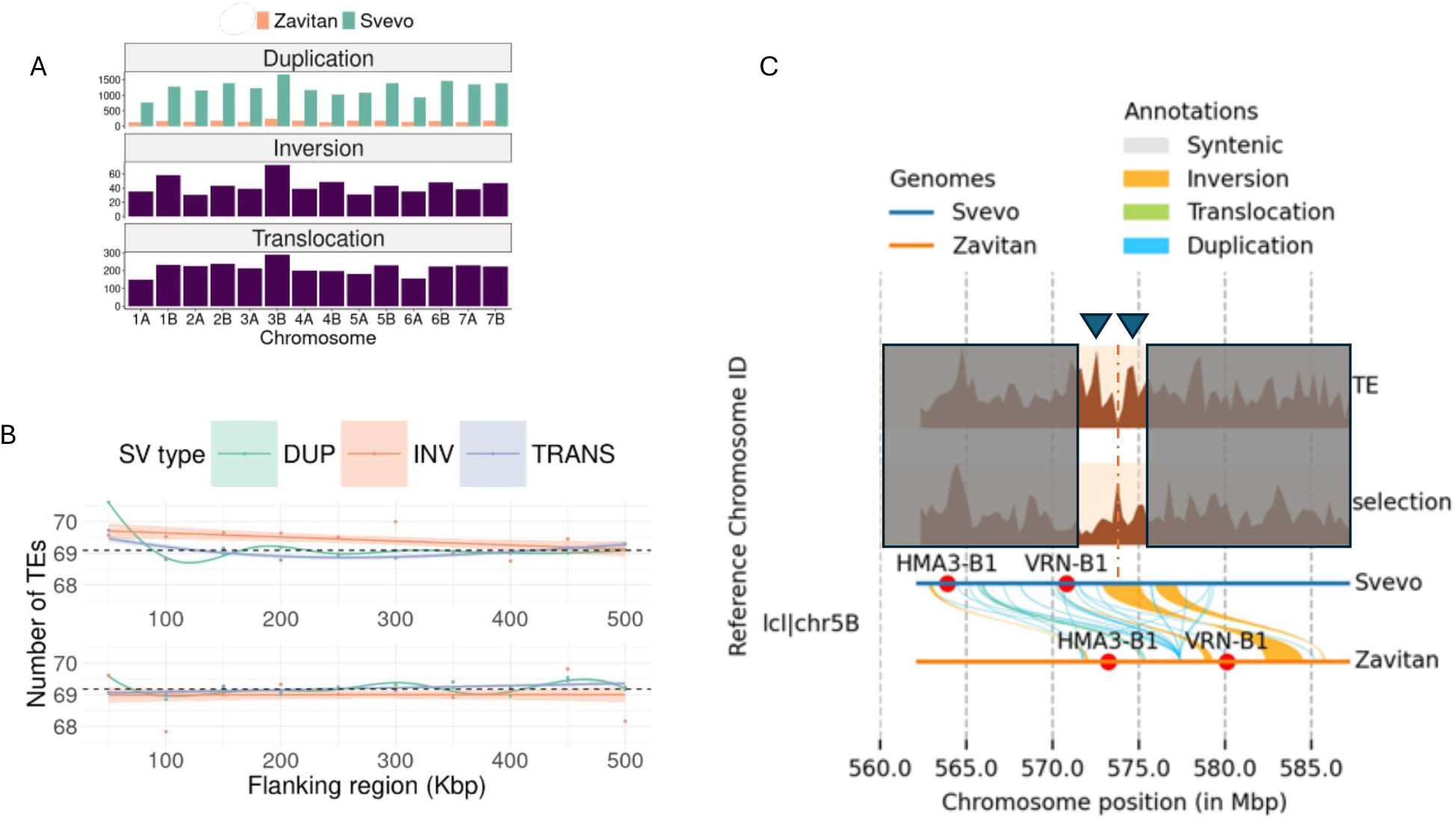
Association between TEs and structural variation (SV). **A)** The distribution of SV types at each chromosome. **B)** Abundance of TEs in the flanking regions around SVs. The number of TEs was calculated within flanking regions (both sides) of each SV using 50Kbp windows. Dots represent the average number of TEs per window for each SV type and marked is the trend line. The upper plot shows the TE abundance around SVs in the Svevo genome and at the bottom is the abundance in the Zavitan. The base line of neutral abundance of TEs along the genome is indicated in horizontal dashed line. **C)** A 20Mbp genomic region containing domestication-related genes on chromosome 5B which also contains several SVs. In the lower panel indicated the alignment between the two reference genomes, Zavitan (bottom) and Svevo, the different SVs colored by type, and the genes in this region in red dot. In the upper panel indicate the TE abundance, and footprints of selection based on Svevo coordinates.

To examine whether SVs are preferentially located in regions enriched with TEs, we quantified the abundance of *Gypsy, Copia*, and *CACTA* elements within the flanking regions of each SV using 50Kbp windows of both sides. As a baseline, TE abundance was estimated from a random set of genomic regions located 1Mbp away from SVs. Overall, TE abundance varied between SV types, with inversions showing consistently higher TE density in the flanking regions compared to duplications and translocations (Fig. 3b). In both Svevo and Zavitan, TEs density around inversions remained above the neutral baseline, indicating localized TE enrichment near inversion breakpoints. Duplications displayed a moderate TE excess in the closest 50Kbp window, which decreased rapidly with distance, and the profile of TEs around translocations was close to neutral. Interestingly, differences in the profile of TEs at the flanking regions were observed between the wild and domesticated reference genomes. The Svevo profile was characterized with sharper TEs enrichment near inversions and stronger divergence between SV types. In contrast, the wild Zavitan profile was smoother with mild differences along the flanking regions, suggesting a potential domestication effect on the abundance of TEs also around SVs.

To investigate whether SVs colocalize with dTE-enriched genomic regions under domestication selection, we generated a null distribution using a permutation test. We calculated the probability of random overlap by simulating genomic regions of matched sizes 10,000 times. The observed overlap between SVs and domestication-related regions was significantly higher than expected by chance (Z = 6.846, p = 0.0001; Fig. S18-19). These results indicate a significant association between domestication selection, TE hotspots, and SVs.

## Discussion

### Domesticated wheat has more TEs than in the wild

Domestication marks a turning point in the history of humanity and in the genomic composition of domesticated wheat. Transposable elements (TEs) comprise a large fraction of the wheat genome, similarly to other major crop species cultivated around the world. Despite their abundance in the genome, TE insertions are generally deleterious because they frequently disrupt gene function when integrated into coding or regulatory regions or by inducing ectopic recombination and structural modifications (Wicker et al. 2018). Among wild emmer wheat (WEW) accessions, we observed that TE polymorphisms largely segregate at low frequencies, reflecting strong purifying selection against their accumulation (Fig. S10). This pattern is consistent with reports in *Arabidopsis*, where nearly all TE insertions within or near genes exert deleterious effects and thus efficiently purged by natural selection (Quadrana et al. 2016). Purifying selection is the primary mechanism removing deleterious TEs, yet it appears to act less efficiently among domesticated wheat, which has experienced restricted admixture compared with WEW and relaxed natural selection under cultivation, thus higher abundance of TEs was observed among domesticated compared to wild accessions (Fig. 1-2). The shift from a wild species to a domesticated crop has reshaped the genomic landscape through demographic bottlenecks and strong positive selection, both of which influence TE dynamics. The reduction in effective population size enhances the genetic drift and weakens purifying selection, allowing mildly deleterious TEs to persist (Wright et al. 2005; Hufford et al. 2012). Long terminal repeat (LTR) retrotransposons dominate the wheat genome, yet domestication did not affect all LTR families equally (Fig. S8). Recombination is naturally suppressed in pericentromeric regions, reducing the efficiency of selection and TE purging. Accordingly, *Gypsy* elements, which preferentially accumulate in these regions, show similar general abundance among wild and domesticated genomes. In contrast, *Copia* elements are more abundant along chromosomal recombining regions where they are purged more efficiently by purifying selection (Fig. 1f, Fig. 2c). Specific regions of suppressed recombination near domestication-related genes, retain dense clusters of TEs, forming hotspots where domestication genes and TEs co-localize (Fig. 1c, Fig. 2c, Fig. 2d). Previous studies on retrotransposon insertion polymorphisms supported their higher abundance among domesticated accessions. For example, specific insertions unique to domesticated wheat (*Jeli–5S* elements) indicate recent mobilization of the *Jeli* family within the last ∼10,000 years (Civán et al., 2013). Thus, despite demographic bottlenecks, TE activity remains an important driver of genomic variability in domesticated wheat. In some cases, beneficial TE insertions that contributed to domestication were already present in wild progenitors and subsequently were fixed by selection. A well-known example is the *Hopscotch* retrotransposon insertion upstream of the *teosinte branched1 (tb1)* gene in maize, which underlies the reduced branching phenotype characteristic of domesticated maize. Molecular dating shows that this insertion predates domestication by over 10,000 years, demonstrating that selection acted on standing TE variation rather than novel insertions (Studer et al. 2011). In wheat, one of the key domestication genes is the *btr1* located on chromosome 3B, a major determinant of the non-brittle rachis phenotype that distinguishes domesticated from wild forms (Harlan et al. 1973). A 4Kbp insertion at the end of the *btr1* gene was previously shown to cause a loss-of-function mutation contributing to the loss of seed shattering phenotype (Avni et al. 2017). A recent long reads assembly of the Langdon variety indicated that this insertion is much longer and spans over 20Kbp (Chen et al. 2025, Fig. S20). Here, both domesticated and wild assemblies were generated from short reads which are less efficient in resolving repetitive regions. These assemblies are likely underrepresented with TEs, but the identified trends were supported using different methodologies including reference-free. Our results confirm that the *btr1* region was under strong domestication selection and that is enriched for fixed TEs among domesticated wheat (Fig. 2d, Fig. S20). Specifically, the insertion consists of *Gypsy* element, which was identified in both Svevo and Langdon genomes, and that this TE insertion disrupted the *BTR1* function leading to the non-shattering phenotype (Avni et al. 2017). This phenomenon is not unique to the *btr1* gene and additional TE hotspots were identified near other domestication-related genes, including the *Q* gene, dormancy-associated loci and vernalization gene (Fig. 1c, Fig. 2c, Table S3). Although no direct functional disruption was detected in these cases so far, the clustering of TEs in selected regions suggests that TE insertions and selection together shaped key phenotypic transitions during wheat domestication. Interestingly, the sequence divergence of these TEs was largely low indicating the insertion was quite recent, presumably during the Pleistocene-Holocene transition.

### Domestication from standing genetic variation

Our results indicate that TEs are more abundant among domesticated wheat (DEW and DW) than in the WEW populations. Similar signatures of TE sequence divergence were observed in both wild and domesticated reference genomes, indicating that major TE proliferation in wheat occurred before domestication (Fig. 1b, Fig. S3). The identified bursts coincide with major climatic turnovers during the transition from the Pliocene to the Pleistocene, the Mid-Pleistocene transition (MPT) and the Marine Isotope Stage 11 (MIS11); a period characterized by exceptionally high temperatures and environmental instability in the Levant (Hu et al. 2024; bartov et al., 2002; Bar-Matthews et al., 1999). Environmental stress including temperature fluctuations, drought, or radiation can trigger episodic TE mobilization and proliferation thus increasing the standing genomic variation (Capy et al., 2000; Quadrana et al. 2016; Rech et al., 2019). Particularly interesting, is the Pleistocene–Holocene boundary period which was characterized by a rapid warming, changing precipitation patterns, and heightened seasonality (Bousios and Gaut, 2016; Horváth et al., 2017). Respectively, a contemporary proliferation event was also observed in both wild and domesticated wheat genomes and supported by low divergence TEs (Fig. 1b, Fig. S3). Interestingly, some of these low diverging TEs were detected around and within domestication related genes, thus suggesting that some of the insertions occurred close to domestication (Fig. 1c).

Previous experimental studies support these climate-linked effects. For example, in *Arabidopsis*, TE transposition is modulated by temperature and precipitation variables, with specific families showing temperature-dependent activation when silencing is impaired Stuart et al., 2016; Quadrana et al. 2016). Similarly, in rice, cold stress induces widespread activation of LTR retrotransposons while suppressing overall gene expression (Yasuda et al., 2013; Negi et al., 2016). Such stress-induced TE mobilization increases mutational input and the emergence of large-effect alleles, potentially accelerating adaptation or the generation of deleterious variants which may be selected to suit specific cultivation needs (Makarevitch et al., 2015). Following the detected ancient proliferation bursts in wheat (Fig. 1b; Charles et al. 2008), purifying selection likely reduced TE abundance in a gradual extended process, removing deleterious insertions while maintaining phenotypic diversity. The elevated standing genetic variation induced by TEs activity may have generated intermediate phenotypes as in the case of the *btr1* gene leading to reduced seed shattering. Early farmers, that were gathering these plants could have identify and select such variants, fixing them through repeated sowing and followed by hybridization between partially domesticated lineages (Purugganan and Fuller 2009, Lev-Mirom et al. accepted).

We therefore propose a “burst-and-select” domestication model for wheat in which environmental stress prior to domestication triggered TE proliferation, expanding phenotypic diversity beyond the variation observed today in natural populations. Thus, reduced shattering may have been more prevalent in WEW populations 10-20Kya than it is today. Subsequent purifying selection filtered gradually such deleterious phenotypes in the wild, while domestication traits were identified and maintained by humans. This model highlights the interaction between environment and genome response in enhancing the potential for domestication. This process may have been similar also in other species facilitating the domestication of the Neolithic founder species.

### TEs reshaped the domesticated wheat genome

The interplay between TEs, recombination, and structural variation (SV) is fundamental in shaping genome architecture and evolution. TEs are major generators of SVs and their genomic distribution closely mirrors local recombination rates (Wicker et el. 2018, Jedlicka et al. 2020). We show that TEs are enriched around SVs and decline with distance (Fig. 3b). Moreover, domestication-specific TEs (dTEs) tend to cluster around domestication-related genes (Fig. 2c–d). Although our SV detection is based on reference genome comparisons and cannot determine whether these SVs arose during domestication or represent ancestral polymorphisms, their overlap with domestication regions under selection, and the gradual decline of TEs with distance suggests a potential link between TEs, SVs, and the phenotypic diversity in the cultivated gene pool. This pattern supports the role of selection in shaping TE persistence and, consequently, evolutionary trajectories (Feschotte & Pritham, 2007; Mézard, 2017).

In wheat and its relative species, TE activity is a major source of gene presence/absence variation (PAV), reflecting recurrent TE-mediated duplications and deletions (Avni et al. 2017, Wicker et al. 2018). Locus turnover rates within the *Triticum–Aegilops* complex correlate strongly with recombination activity (Dvorak et al., 2018). Recent TE insertions are concentrated in gene-rich, recombinogenic chromosomal regions, highlighting the spatial coupling between recombination, TE dynamics, and functional gene content (Wicker et al., 2018; Keidar-Friedman et al., 2018). Both TEs and SVs can generate large-effect alleles rapidly favored by selection (e.g. *btr1*) and can give rise to novel regulatory elements or proteins. Positive selection on advantageous TE insertions or SVs produces distinct genomic footprints, reflected as peaks of differentiation between wild and domesticated accessions (Fig. 2c–d). TE clusters may also promote ectopic recombination, further contributing to SV formation. However, the precise mechanisms linking TE activity and SV induction during domestication remain largely elucive. The observed association between TEs, recombination, and selection around domestication genes is not unique to wheat. For example, a TE-associated selective sweeps have also been detected near the *Y1* locus in maize, and next to the *waxy* gene in rice (Palaisa et al. 2004; Olsen et al. 2006).

We highlight the interaction between the wheat genome and the environment in the context of domestication which reshaped not only plant phenotypes but also genomic architecture fostering new genetic variation and enhancing population differentiation. While many such variants were likely purged by purifying selection in wild populations, their transient abundance during early Holocene climates may have provided the raw material that early farmers selected upon. Assuming these phenotypes were more prevalent 10–20K years ago, domestication likely stemmed from early farmers who capitalized on the advantageous phenotypes they observed.

## Materials and Methods

### Annotation of TEs and estimating bursts events over time

To ensure accurate annotation and maximize coverage, a custom Transposable Element (TE) library was constructed for the Triticum turgidum ssp. durum (Svevo v1; Maccaferri et al., 2019) and wild emmer wheat (T. dicoccoides Zavitan v1.0; Avni et al., 2017) reference genomes. We combined de novo predictions from both genomes with the public TREP database (v19; Wicker et al., 2002). Initial candidate LTR retrotransposons were identified using LTR_FINDER_parallel v1.3 (Ou & Jiang, 2019) and LTR_harvest v1.6.6 (Ellinghaus et al., 2008). These candidates were filtered and curated using LTR_retriever v3.0.4 (Ou & Jiang, 2018) to ensure high-confidence structural annotation. This curated *de novo* library was merged with the TREP database, and the combined dataset was clustered using CD-HIT-EST v4.8.1 (Fu et al., 2012) at a 90% sequence identity threshold to eliminate redundancy, resulting in a final comprehensive consensus database.

Using this custom consensus library, we performed a standard annotation of TE families across both reference genomes using RepeatMasker v4.1.8 (Smit and Hubley, 2025) with default parameters. Following the initial annotation, we performed a focused abundance profile analysis by retaining only the LTR/Gypsy, LTR/Copia, and TIR/CACTA superfamilies, which represent the most abundant and widely distributed TE classes in wheat genomes.

To reconstruct the evolutionary history of these elements, we employed two complementary approaches to estimate insertion times, a consensus-based age estimation and pair-wise divergence estimates. For the consensus-based estimation, we calculated the percentage of nucleotide divergence (*D*) of each annotated TE copy from its consensus sequence directly from the RepeatMasker output. This divergence score is used as a proxy for the accumulation of neutral mutations since insertion. We converted this divergence score into an estimated age in years using the standard formula (Bowen and McDonald, 2001; Jedlicka et al., 2020):

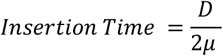

Where *D* is the percentage of divergence and *µ* is the neutral substitution rate of 1.67×10^-8^ substitutions per site per year (Cantu et al., 2010). This approach provided the overall age distribution of TE families relative to their ancestors.

For the pair-wise divergence estimates highlighting recent proliferation dynamics, we calculated the divergence of each TE sequence against its closest hit within the same genome. Significant hits were identified using BLASTN v2.14.1 (Altschul at el. 1990), and the level of divergence between pairs was used as an indication for the time since insertion, thus providing a targeted view on recent insertion events. Gene coordinates in the Svevo reference genome were extracted directly from the annotation file. For validation of the *BTR* genes we also aligning the barley *BTR1* and *BTR2* sequences (Sato et al. 2021) to the Svevo genome using GMAP version 2025-04-19 (Wu and Watanabe 2005).

### Tetraploid wheat diversity panel

A diversity panel for tetraploid wheat was recently developed to represent the main three groups: wild emmer wheat (*T. turgidum* ssp. *dicoccoides*; WEW), domesticated emmer wheat (*T. turgidum* ssp. *dicoccum*; DEW), and durum wheat (*T. turgidum* ssp. *durum*; DW) (Klymiuk et al. 2023; Lev-Mirom et al. accepted). Briefly, the collection was developed from a larger dataset of over 500 accessions which was genotyped using the 90K SNP chip and analyzed to identify a representative core collection that captures most of the diversity among tetraploid wheat groups. The final core collection which was used here is comprised of 70 WEW from across the distribution range of the species, 44 DEW and 45 DW landraces from all over the globe. Additional elite cultivars that were included in the original core collection were excluded; thus, a total of 159 accessions were included in this study.

Genotype data called from whole-genome sequence data was available for the entire core collection (Lev-Mirom et al. accepted). Briefly, the core collection was whole-genome sequenced using Illumina short-read technology for each accession. Raw sequences were cleaned with the fastp software v0.20 (Chen et al. 2018), aligned with BWA-MEM2 v2.2.1 (Vasimuddin et al. 2019) to the Svevo reference genome v.1 (Maccaferri et al. 2019), and variants were called using GATK v4.1.9 (McKenna et al. 2010) following the best practice recommendations. Variants were extensively filtered to remove low quality calls, heterozygote sites, variants with minor allele frequency lower than 5% and maximum of 10% missing data. The final data set included 6,747,640 variants called across all 159 accessions.

### Assignment of accessions to populations

To explore the classification of populations and the assignment of accessions to genetic clusters, genotype data was pruned to reduce bias introduced by linkage disequilibrium (LD) between SNPs. Pruning was conducted with PLINK v1.9 (Purcell et al. 2007) at a minimum LD cut-off of *r*^*2*^ = 0.2 which was calculated in windows of 100Kbp and step size of 10Kbp. Following pruning, 2,656,229 SNPs were kept for a population structure analysis as implemented in fastSTRUCTURE v1.0 (Raj et al. 2014). To identify potential mis-assignment and admixture between wild and domesticated wheat, the number of clusters was set to K = 2 with 5 rounds of cross validation. Accessions with less than 50% assignment to their expected group as defined in the meta-data information were removed. Next, the same procedure was conducted only for the domesticated accessions with K = 3 to resolve potential misassignment among DEW and DW. Population stratification among tetraploid wheat groups was also explored using a principal component analysis (PCA) as implemented in PLINK v1.9 using the same pruned dataset. The top two principal components (PC1 and PC2) were used to generate a visualization after coloring accessions by their expected group.

### Population genomics and identification of footprints of selection

Population statistics including average nucleotide diversity (π), Tajima’s D, and population differentiation (*F*_*ST*_) were calculated in non-overlapping windows of 50Kbp using VCFtools v0.1.16 (Danecek et al. 2011). Reduction in diversity during domestication was calculated in each window by dividing the domesticated group score by the wild group score (πD/ π_W_).

Recombination rates (ρ) were estimated using LDhat v2.2 (Auton and McVean 2007). Before running LDhat, SNPs were thinned to one SNP every 1000bp. Variants were divided into batches of 5000 SNPs with an overlap of 500 SNPs between consecutive batches. For each batch, recombination rate estimation was performed with the *interval* program in LDhat using a pre-computed likelihood lookup table generated for a population-scaled mutation rate of *θ* = 0.001 and for the corresponding sample size. Markov chain Monte Carlo (MCMC) simulations were run for 10,000,000 iterations with a block penalty of 5 to control rate variation between adjacent sites, and posterior samples were collected every 500 iterations. The first 15,000 iterations were discarded as burn-in period. Posterior summaries of *ρ* were obtained with the *stat* program in LDhat, and *ρ* estimates were averaged across non-overlapping 250Kbp windows. To identify regions where recombination was reduced following domestication, we computed the ratio of recombination rate in domesticated to wild accessions (ρ_*D*_/ρ_W_).

To detect genomic regions that were targeted by selection during the transition from wild to a domesticate, we integrated the signal obtained from the calculation of π_D_/π_W_, F_ST_, Tajima’s D and ρ_*D*_/ρ_W_ at each window into a single score. The scores of each statistic were transformed into a rank-based log_10_(*P*-value) in accordance with the direction of the score, thus low Tajima’s D, ρ and π, were transformed using the formula:

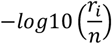

where *r*_*i*_ is the rank of a window and *n* is the total number of windows. For population differentiation index (*F*_*ST*_), high values were ranked as the top scores and the transformation was conducted with the formula:

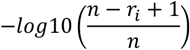

The resulting standardized scores were then combined using the Mahalanobis distance method (Lotterhos et al. 2017) into a composite score for each 50Kbp window. Finally, the top 5% of windows were considered as candidate windows that were targeted by selection during domestication.

### *De novo* TE quantification

To quantify the abundance of TEs along the genome of each accession in the diversity panel, whole genome sequence data was analyzed using the dnaPipeTE software v1.3.1 (Goubert 2023). This pipeline enables to find, annotate and quantify repetitive elements in a representative fraction of whole genome sequencing data and therefore is specifically suitable for low coverage sequencing data. The pipeline performs a *de novo* assembly of the repetitive sequence without relying on a reference genome thus avoiding reference bias in the analysis. The *Triticeae* Repeat Sequence Database (Wicker et al. 2002) was used as the TE library in the analysis to search and annotate repeats.

To avoid a sampling bias in the estimates of repetitive elements frequencies due to variation in coverage between samples, a subset of reads that properly represent the genome composition of each genotype was randomly sampled and used in the analysis. To determine the minimum number of reads required to properly represent the genome composition, the analysis was performed with varying subsets of reads sampled from a single individual. The number of reads tested ranged between 500,000 to 20,000,000 and the proportion of repetitive elements identified was evaluated at each round. Once the proportion of the different types of repetitive elements remained unchanged along exceeding number of reads, sampling was considered saturated. The minimum number of reads to reach saturation was at 10,000,000 reads which was fixed as the number of reads sampled across all accessions. The proportion of TE-derived base pairs per accession was calculated as the number of bases aligned to a TE sequence divided by total sampled bases.

### Identification of TE insertions

To identify non-reference TE insertions, we used INSurVeyor v1.0 (Rajaby et al 2023) which can detect sequence insertions from discordant read pairs and split-read alignments in short-read data. All accessions were analyzed using alignments data to the Zavitan reference genome because the wild reference is ancestral to the domesticated type and the analysis identifies insertions with respect to the reference. Indeed, running the same analysis with the Svevo as reference yielded less than half of the number of insertions as with Zavitan. To restrict the analyses only for TEs insertions, we filtered all occurrences that were not annotated as TEs based on the TREP database. To consolidate insertions identified in each accession into one comparative table, we employed the SurVClusterer v1.0 (https://github.com/Mesh89/SurVClusterer) to merge all TE insertions based on genomic coordinates and identity. This approach enabled to obtain a unified non-redundant TE insertion table across accessions. Finally, the TE insertions occurrence table was used to generate a site-frequency spectrum (SFS) for domesticated and wild groups.

### Generating a TE presence/absence variation matrix across all accessions

Sequence data from each accession were aligned to the Svevo and to Zavitan reference genomes using BWA-MEM2 (Vasimuddin et al. 2019). To identify TEs that are associated with wheat domestication, we summarized the genomic positions of all TEs annotated in each reference genome using RepeatMasker into a BED file format. For each accession, the alignment file (bam) was integrated with the TEs positions in the bed files to extract the sequencing depth and coverage using bamdst v1.1.0 (https://github.com/shiquan/bamdst). The resulting coverage and depth scores were organized into a table which include all accessions and the corresponding percentage of each TE sequence coverage. This table was then converted into a presence/absence variation (PAV) matrix, where TEs were considered present if coverage exceeded 75% of the sequence length, absent if no reads were aligned (zero coverage), and unknown otherwise. This conservative approach enables to reduce the rate of false identifications of TEs with a cost of missing many potential TEs, thus reducing the number of TEs analyzed. Nevertheless, due to the abundance of TEs along the genome we were more concerned about increasing the number of falsely identified TEs which may inflate the overall number of TEs and dilute the signal. In addition, the entire procedure was also performed using the Zavitan genome as reference to guarantee results are not reference-dependent (Fig. S12). For all subsequent analyses, we used the PAV matrix obtained using the Svevo as reference due to higher coverage of TEs.

### Identification of domestication TEs

To identify TEs that differentiate between wild and domesticated wheat a principal component analysis (PCA) was conducted using the PAV matrix. Based on the PCA, we extracted the loading score for each TE from PC1 which is indicative for the relative contribution of each TE to the overall split between wild and domesticated wheat. Negative loading scores were attributed to TEs that are found in high frequencies among domesticated accessions (dTEs) but largely missing in the wild. In contrast, positive loading scores were considered wild-related TEs (wTEs). Extreme scores at the tails of the loading distribution have the strongest impact on the differentiation, thus the most negative loading for dTEs are insertions that were fixed among domesticated wheat. The lowest 1% TEs were extracted and counted at windows of 50Kbp for further comparison with other genome scan analyses. Visualization in a circular Manhattan plot was generated with the CMplot package v4.5.1 in R (Yin et al. 2021).

### Structural variation analysis

To identify structural variation between wild and domesticated wheat we compared the Svevo and Zavitan reference genomes using SyRI v1.6 (Goel et al. 2019) and MUMmer v4.0.1 (Marçais et al. 2018). Analyses were conducted for each chromosome separately to maintain a manageable computation load with a cost of missing inter-chromosomal translocations. Alignments between genomes were conducted using the genome alignment function with the ‘maxmatch’ option to guarantee all anchor matches are used. The exact minimum match was set to 50bp which were clustered if they spanned at least 100bp with a maximum of 500bp gap permitted between clusters. The alignment files were then filtered for minimum of 90% identity and at least 100bp length to remove spurious alignments. Parameters in MUMmer were set in accordance with the SyRI pipeline recommendations. Structural variant (SV) calling was conducted using the alignment files and included inversions, translocations, duplications, and deletions, where the Zavitan genome was used as the query sequence. To minimize false-positive SVs detection arising from TEs or alignment artifacts, we applied an additional filtering step based on TE content. Coordinates of all annotated TEs from both the Svevo and Zavitan reference genomes were aligned to the identified SV coordinates. For each SV, we calculated the proportion of base pairs overlapping annotated TE regions using GRanges (Lawrence et al. 2013). SVs with more than 95% overlap with TEs were considered TE-derived and were excluded from the analysis. Lower thresholds (down to 50%) were also tested and the trend was consistent.

To test for potential overlaps between SVs, dTE hotspots, and regions under selection we used the GRanges and visualized using the eulerr package (Larsson and Gustafsson 2018). A permutation test was conducted using the permTest function from the regioneR package (Bernat G et al. 2016) to evaluate if the number of overlaps significantly deviate from random. A null distribution was generated using 10,000 iterations, where in each iteration a random region of the same length as the tested SV was extracted. Finally, the number of random overlaps was compared to the number of true overlaps between SV and regions of selection that are enriched with dTEs (Fig. S19).

## Acknowledgements

We thank Evgenii Potapenko for his valuable advice on the computational methodology throughout the study. This study was supported by the Israel Science Foundation (ISF) grant ISF-1154/19 (SH).

